# Heat production in a feeding matrix formed on carrion by communally breeding beetles

**DOI:** 10.1101/854349

**Authors:** Szymon Matuszewski, Anna Mądra-Bielewicz

## Abstract

Insects regulate their body temperature mostly behaviourally, by changing posture or microhabitat. These strategies may be ineffective in some habitats, for example on carrion. Carrion beetles create a feeding matrix by applying to cadaver surface anal or oral exudates. We tested the hypothesis that the matrix, which is formed on carrion by communally breeding beetle *Necrodes littoralis* L. (Silphidae), produces heat that enhances insect fitness. Using thermal imaging we demonstrate that heat produced in the matrix formed on meat by adult or larval beetles is larger than in meat decomposing without insects. Larval beetles regularly warmed up in the matrix. Moreover, by comparing matrix temperature and larval fitness in colonies with and without preparation of meat by adult beetles, we provide evidence that formation of a matrix by adult beetles has deferred thermal effects for larval microhabitat. We found an increase in heat production of the matrix and a decrease in development time and mortality of larvae after adult beetles applied their exudates on meat in the pre-larval phase. Our findings indicate that spreading of exudates over carrion by *Necrodes* larvae, apart from other likely functions (e.g. digesting carrion or promoting growth of beneficial microbes), facilitates thermoregulation. In case of adult beetles, this behaviour brings distinct thermal benefits for their offspring and therefore may be viewed as a new form of indirect parental care with an important thermal component.

---

Temperature is a key component of animal environment. Many species, particularly insects, are more or less dependent on external heat (1, 2). Animals frequently use heat that is already present in the environment, through changing body orientation (e.g. basking), selecting thermally attractive microhabitat or optimising its thermal characteristics (e.g. aggregating) (3-5). Rarely, however, they transform the environment to affect, focus and take advantage of thermogenesis. Some Indo-Pacific megapode birds, e.g. Australian brush-turkey (*Alectura lathami*) construct their incubation mounds from plant litter, where the composting generates heat that is used for egg hatching (6, 7). Some crocodilians make similar incubation mounds, in which heat is produced by decomposing plant material (6, 8, 9). Habitat transformation to facilitate thermogenesis may occur also in carrion insects. Blow flies (Calliphoridae) or carrion beetles (Silphidae) use cadavers mostly for breeding and their larvae are main carrion reducers in some terrestrial environments (10-12). Necrophagous larvae usually feed in aggregations (13), which may have much higher inner temperature than ambient air (by 10-30°C). This effect, called the maggot-mass effect, was originally discovered in blow flies (5, 14-18), but has also been reported for *Necrodes* beetles (Silphidae) (19). The inner heat of these aggregations was hypothesized to derive from microbial activity (20, 21), larval exothermic digestive processes (17) or larval frenetic movements (18, 22). However, there is no evidence to support any of these mechanisms.

Carrion beetles form a feeding matrix on cadavers (also called “a biofilm-like matrix”) by spreading over its surface anal and oral exudates (23). Carrion smearing was originally described in adult burying beetles (Silphidae: *Nicrophorus*) (24). This behaviour was hypothesized to moisturize carrion (24), facilitate digestion (24-26), suppress microbial competitors (23, 27-30), deter insect competitors by reducing carrion-originating attractants (24, 31, 32), support larval aggregation (24) or development (27) or seed mutualistic microbes and transmit them to offspring (25, 33-35). Exudates of adult or larval burying beetles were found to contain antimicrobial compounds (36-40). Moreover, presumptively mutualistic microbes (e.g. *Yarrowia* yeasts) were abundantly identified in carrion beetle guts and the matrix formed on carrion by the beetles (23, 25, 35, 41, 42). These findings indicate that feeding matrix on carrion is a complex microenvironment emerging from interactions between burying beetles, microbes and a putrefying resource. Although cadaver smearing behaviour has not been reported in other carrion beetles, there is indirect evidence suggesting that formation of the matrix is more prevalent among these beetles (36, 41). The matrix probably brings several benefits for the beetles and their scope may vary between the species.

Carrion beetles (Silphidae) are divided into Nicrophorinae (burying beetles), and Silphinae (43). The latter subfamily is more diverse, with necrophagous, predatory and phytophagous species (44, 45). *Necrodes* beetles are members of Silphinae, grouped with *Ptomaphila, Oxelytrum* and *Diamesus* at the base of the subfamily (45-47). *Necrodes* colonizes large vertebrate cadavers, where its larvae feed on carrion tissues and under favourable conditions may reduce them into dry remains (48, 49). Beetles start visiting carrion after it becomes bloated (under summer temperatures usually 4-8 days after death) and after some time many of them (even hundreds) may be present on a cadaver (48, 50-52). Females lay eggs (30-50 per female) in a nearby soil and larvae abundantly colonize carrion during late decomposition (48, 50, 51). Adult beetles are usually absent when larvae colonize the resource. In Central European forests during the summer adult *N. littoralis* were present 3 to 8 days on pig carcasses and usually 5 to 7 days elapsed between the arrival of the first adult beetles and the first larvae (53). Under laboratory conditions, when food and temperature are optimal, adult beetles oviposit within 1-2 days of provisioning them with fresh meat. The egg stage of *N. littoralis* lasts on average 3.4 days at 22°C and 4.9 days at 18°C (Gruszka, personal communication). *Necrodes* larvae feed in aggregations, which form in the warmest place and relocate in response to changes in the heat source location (13). This indicates that heat plays an important role in the formation and maintenance of larval aggregations in *Necrodes* (13). After larvae stop feeding, they pupate in a nearby soil (48). When disturbed, adult *Necrodes* beetles spray defensive anal secretions (frequently mixed with excretions), which have a strong repellent effect against other insects (54, 55) and a significant antibacterial action (36). However, its addition directly to the carrion has not been reported. *Necrodes* beetles, in contrast to burying beetles, colonize large carrion, breed there communally and are considered as species without parental care (44, 45, 50).

We found that *Necrodes littoralis* L. under laboratory conditions forms on meat a feeding matrix, similarly to burying beetles. Because larvae of this species are distinctly heat-oriented, and external heat certainly enhances their fitness, we formulated a hypothesis that the feeding matrix produces heat, which is beneficial for the larvae. Here, we provide evidence that the matrix made from exudates of larval *N. littoralis* has a significantly higher temperature than the matrix formed on meat decomposing without the insects. Beetle larvae were regularly warming up in this microenvironment, supporting the hypothesis that smearing carrion with exudates facilitates thermogenesis and indirectly larval thermoregulation. Because both adult and larval beetles were found to spread their exudates over meat forming the matrix, we wanted to test if the application of exudates by adult beetles affects heat production of the matrix during the larval feeding phase and eventually brings thermal benefits for the larvae. We found an increase in heat of the matrix formed by larvae and a decrease in larval development time and mortality, following the application of exudates by adult beetles. Therefore, spreading the exudates by adult beetles had lagging thermal effects for their offspring’s microenvironment, which supports the hypothesis that *N. littoralis* manifests an indirect thermal parental care.

## Results

### Heat emission in feeding matrix formed on carrion by *Necrodes* beetles

To test if feeding matrix produced by *N. littoralis* generates heat, we monitored with thermal imaging conditions in colonies of adult (A) and larval (L) beetles subsequently feeding on meat (M) (hereafter M+A+L setup) and in colonies of larval beetles only (hereafter M+L setup), using as a reference the equivalent meat setup without the insects (hereafter M setup, Fig. 1). We found that adult and larval beetles spread their exudates over meat to form a greasy, feeding matrix that covers the meat and surrounding soil, the surface of which was enlarging with colony age (Fig. 2). The matrix had a higher temperature than the background (Fig. 3) and beetle larvae were regularly warming up on its surface (Fig. 4).

**Fig. 1.**
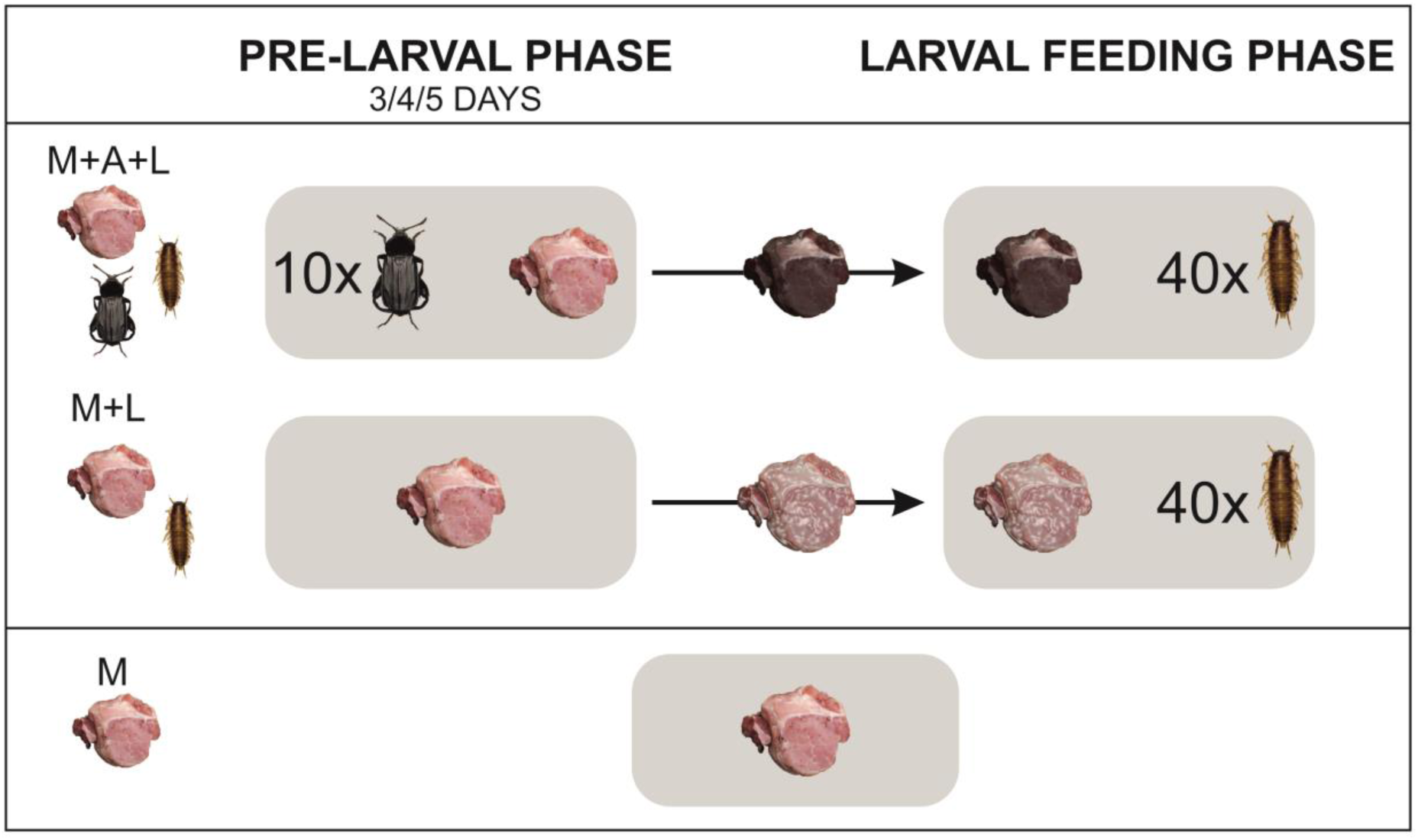
Experimental design to study the heat production of a feeding matrix formed on meat by carrion beetle *Necrodes littoralis* and the development of its larvae in colonies with and without adult beetles presence in the pre-larval phase (respectively M+A+L and M+L setups). Trials with meat only (M setup) were used as the reference.

**Fig. 2.**
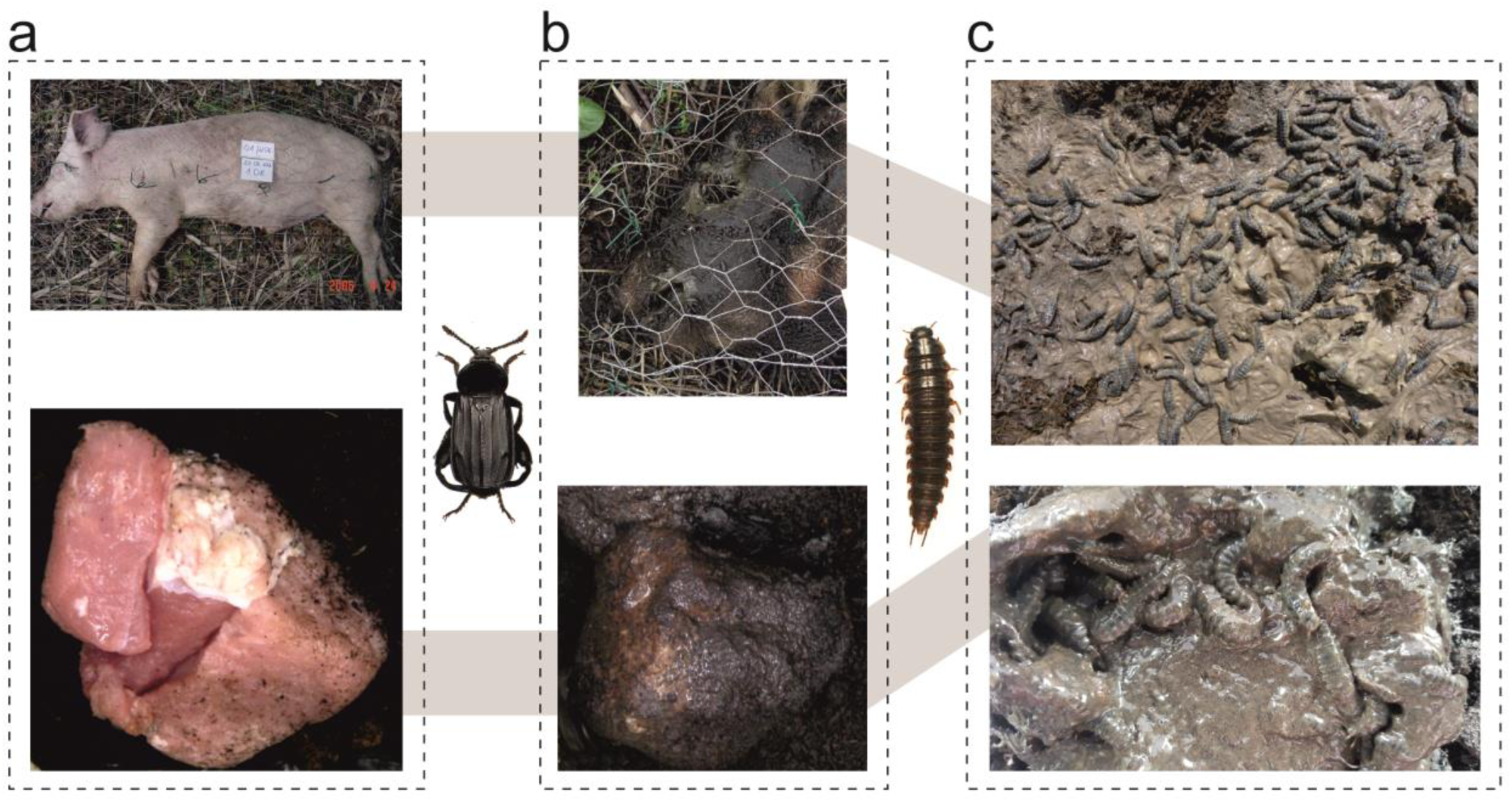
The feeding matrix formed by adult or larval *Necrodes littoralis* on pig cadaver under field conditions (top panel) and on meat in the laboratory (bottom panel). a – fresh resources, b – the matrix formed by adult beetles, c – the matrix formed by larvae.

**Fig. 3.**
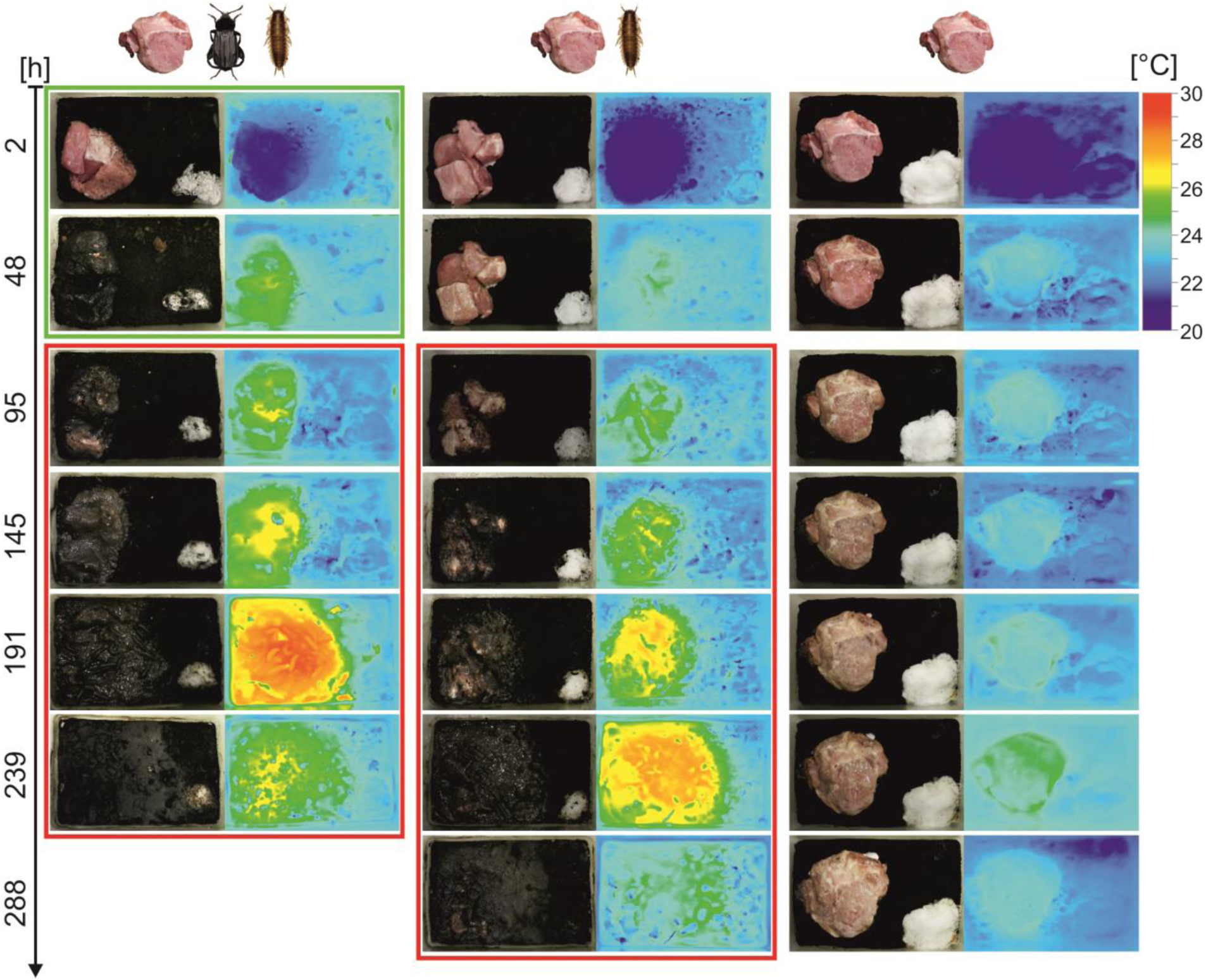
Heat production of the feeding matrix in colonies with adult and larval *Necrodes littoralis* (left column), with larval beetles only (middle column) and without the beetles (right column). The green frame includes pictures of adult beetle colonies, the red frame pictures of larval beetle colonies. Pictures without a frame show meat decomposing without insects.

**Fig. 4.**
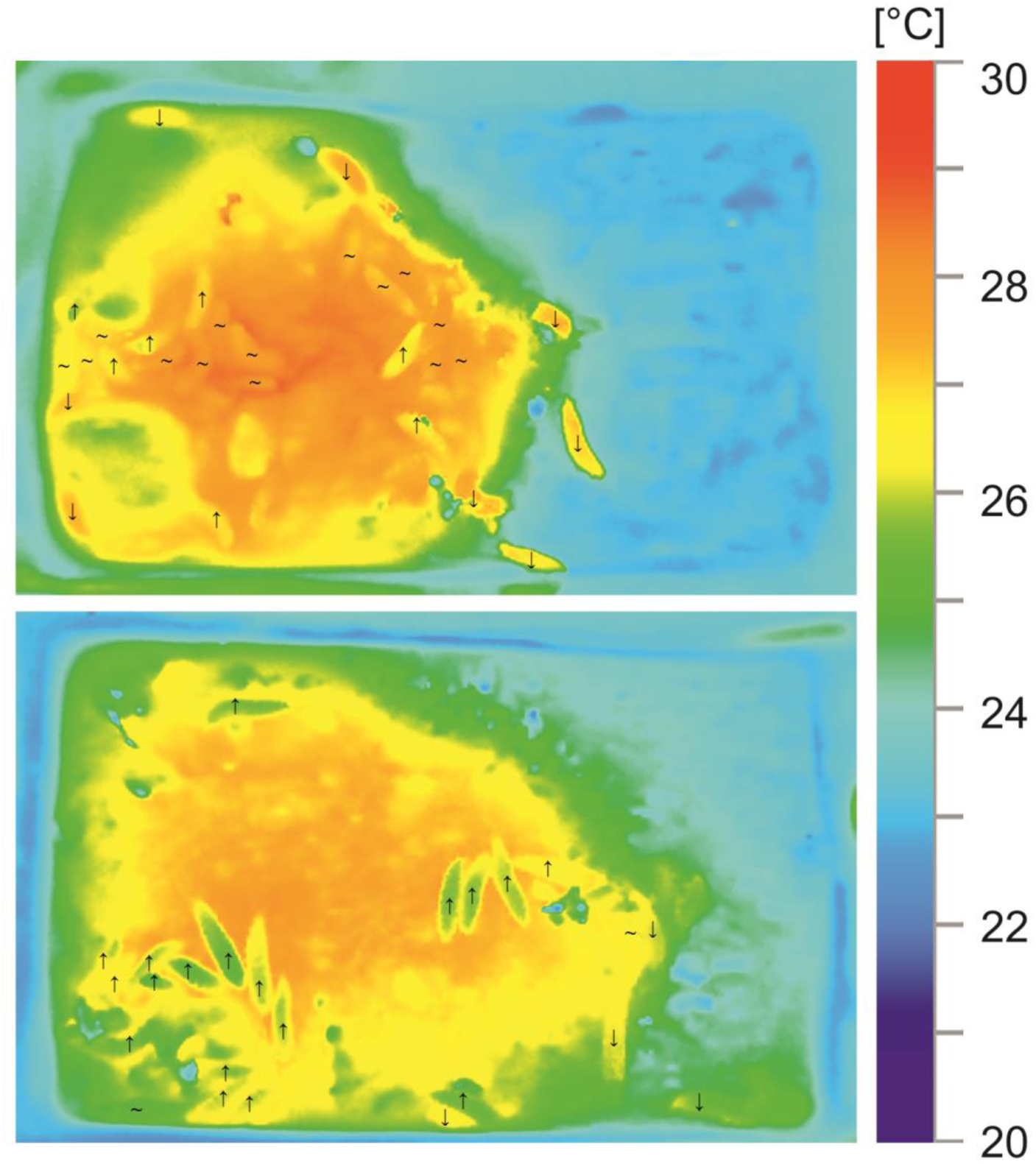
Thermal images of larval *Necrodes littoralis* warming up in the feeding matrix. Images demonstrate that heat of the larvae is a function of the matrix heat. Larvae present on the matrix may be classified into: ‘warming’ (**↑**) and ‘warmed’ (**∼**). Temperature of the ‘warming’ larvae is much lower than temperature of the matrix in their vicinity and temperature of the ‘warmed’ larvae is the same or only slightly lower. There is a clear heat gradient from the outside to the inside in the ‘warming’ larvae, which indicates they are warming up in the matrix. There are also ‘cooling’ (**↓**) larvae, with higher body temperature than in their vicinity and a heat gradient from the inside to the outside. However, they are present only outside of the matrix. Assuming that larvae contribute endothermically to the heat of the matrix, the ‘cooling’ larvae should be present on the surface covered with the matrix. As this is not the case, the images demonstrate that heat is not endothermically generated by the larvae, but is produced in the matrix. Moreover, larvae are warming up while staying on the matrix.

To investigate the effect of environmental temperature on heat emission in a matrix, we compared larval colonies (M+L setup) reared under different temperatures, using as a reference paired containers with meat only (M setup). The comparison revealed that the heat production in the feeding matrix formed by larvae increased with a rearing temperature (*F*_3,35_=59.4, *N*=39, *P*<0.001, Fig. 5a, b in ESM) and was significantly larger than in meat alone at all rearing temperatures, apart from 16°C (*F*_1,35_=87.5, *N*=39, *P*<0.001, Fig. 5b in ESM).

By quantifying the average temperature of a matrix-covered surface (M+A+L and M+L setups) and meat without the beetle-derived matrix (M setup), we found that heat emission in the matrix was significantly larger than in meat alone (*F*_1,27_=122, *N*=30, *P*<0.001, Fig. 6a, b). The temperature of the matrix revealed a steady increase until meat resources were depleted and larvae stopped feeding (Fig. 6a). When adult beetles (M+A+L setup) applied their exudates in the pre-larval phase, the temperature of the matrix was significantly higher in the larval feeding phase, as compared to the colonies of larval beetles only (*F*_1,27_=154.8, *N*=30, *P*<0.001, Fig. 6c). Moreover, heat production in the matrix during the larval feeding phase became larger, when we extended the pre-larval phase (*F*_2,27_=24, *N*=30, *P*<0.001, Fig. 6a, c).

**Fig. 6.**
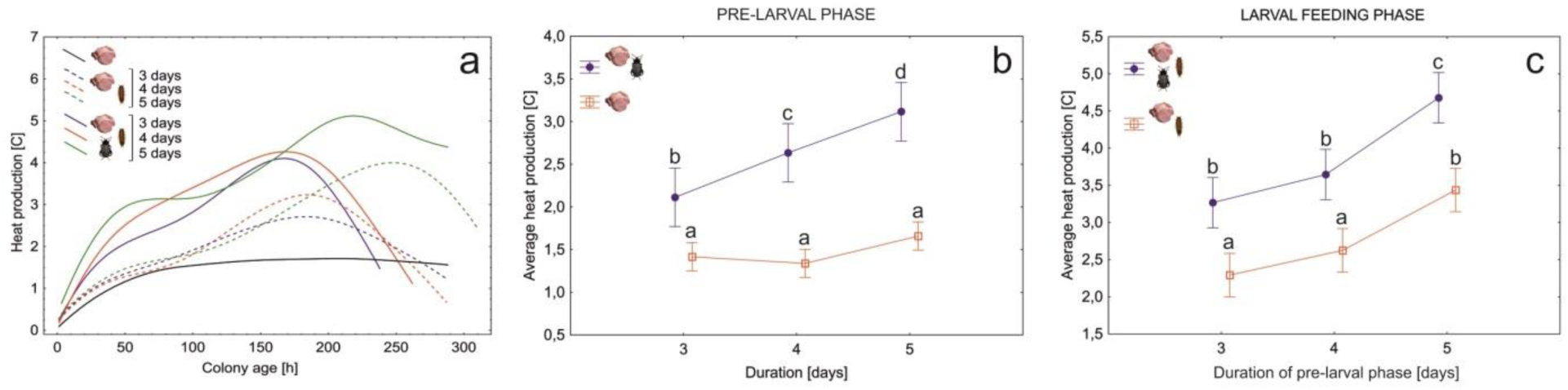
Longitudinal thermal profiles (a), differences in the average heat production of the feeding matrix during the pre-larval phase (b) and the larval feeding phase (c) between *Necrodes littoralis* colonies differing in the presence of adult beetles and in the length of the pre-larval phase. All colonies were reared at constant 23°C. As a no-beetle reference in (a), we used results for 23°C from the environmental temperature experiment. Thermal profiles were fitted to the data using the distance-weighted least-squares smoothing procedure. Symbols – means, whiskers – 95% confidence intervals, different letters denote significant differences in pairwise comparisons.

### Feeding matrix, larval fitness and parental care in *Necrodes* beetles

To test if spreading the exudates over meat by adult beetles affects the fitness of larvae, we compared development times of larvae and pupae, larval mortality and mass of postfeeding larvae between colonies with and without the presence of adult beetles in the pre-larval phase (M+A+L and M+L setups). Larvae developed significantly shorter after adult beetles prepared the meat (*F*_1,27_=15, *N*=30, *P*<0.001, Fig. 7a), differences in the pupal development time were insignificant (*F*_1,27_=0.8, *N*=30, *P*=0.39, Fig. 7b). Mortality of larvae was significantly lower in colonies with meat prepared by adult beetles (*F*_1,27_=8.3, *N*=30, *P*<0.01), differences between the setups were the largest under the shortest pre-larval phase (Fig. 8a). An increase in the duration of the pre-larval phase resulted in higher larval mortality (*F*_1,27_=3.5, *N*=30, *P*=0.045, Fig. 8a). Application of exudates by adult beetles revealed no significant effect on the mass of postfeeding larvae (*F*_1,27_=2.4, *N*=30, *P*=0.13, Fig. 8b).

**Fig. 7.**
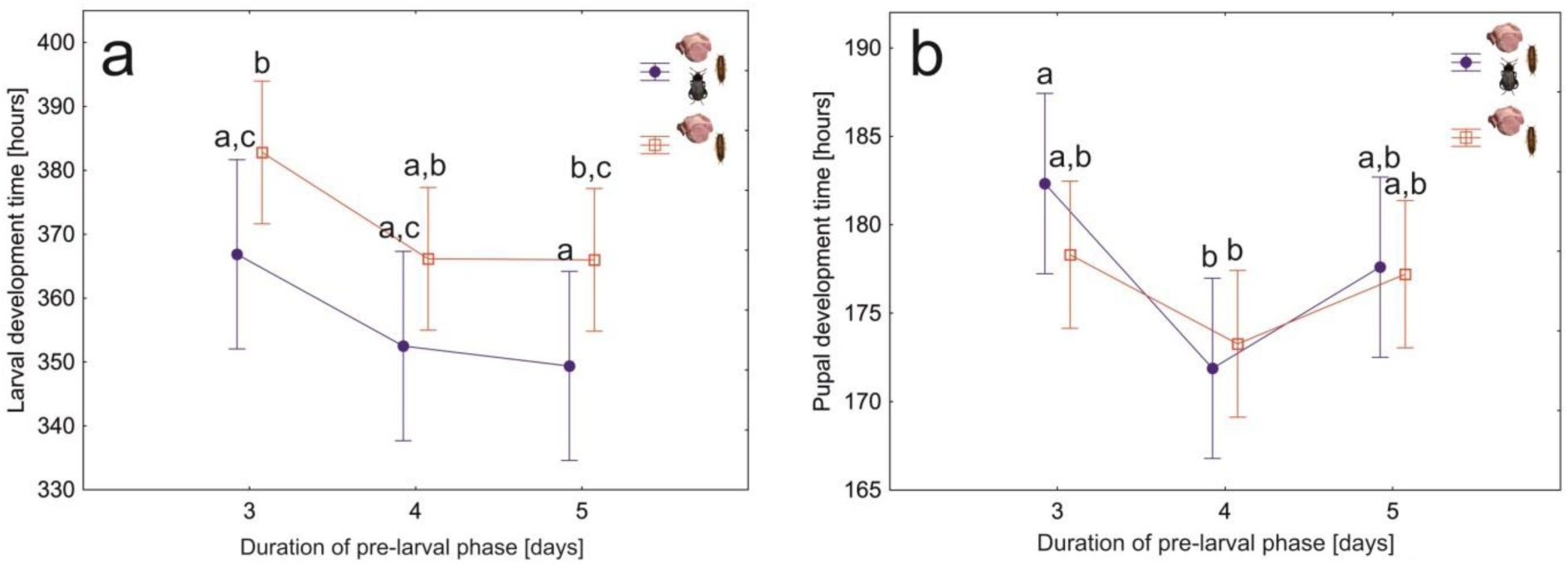
Average larval (a) and pupal (b) development times for *Necrodes littoralis* colonies differing in the presence of adult beetles during the pre-larval phase and in the duration of this phase. Larval development times include feeding (on meat) and postfeeding (in Petri dishes) phases. Symbols – means, whiskers – 95% confidence intervals, different letters denote significant differences in pairwise comparisons.

**Fig. 8.**
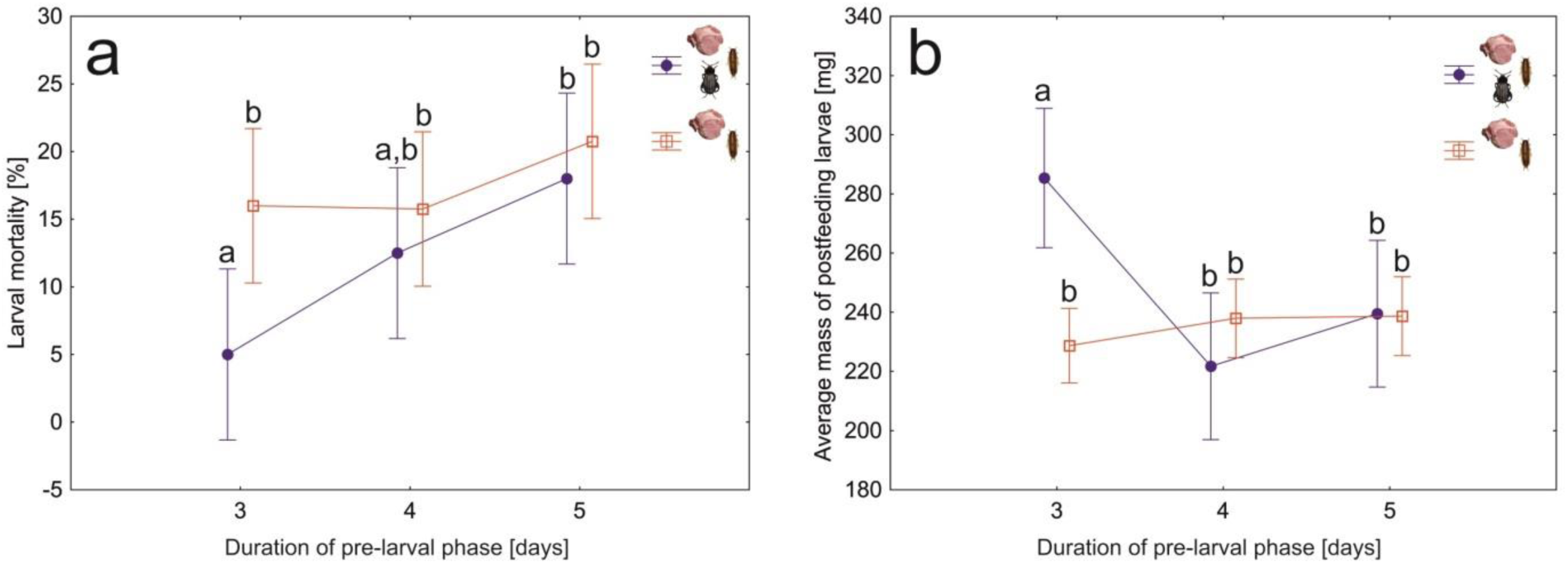
Larval mortality (a) and average (per larva) mass of postfeeding larvae (b) for *Necrodes littoralis* colonies differing in the presence of adult beetles during the pre-larval phase and in the duration of this phase. Symbols – means, whiskers – 95% confidence intervals, different letters denote significant differences in pairwise comparisons.

## Discussion

We demonstrated that *Necrodes* beetles form a feeding matrix on carrion in which heat is produced and beetle larvae warm up while feeding. The heat emission recorded in insect colonies was highly consistent in space and time with the matrix formation (Fig. 3). Since there is no known endothermic mechanism in insect larvae (2, 5), we rejected a hypothesis that the larvae contributed endothermically to the heat increase. Even if some unknown mechanism operated in our colonies, they consisted of only 40 larvae, so the mechanism would probably be not effective in raising and maintaining the temperature over the entire matrix-covered surface. Moreover, our close-up thermal images (Fig. 4) demonstrated that the heat of the larvae was a function of the heat of the matrix and not vice versa. Therefore, we interpret our findings as evidence that the heat is not endothermically generated by the larvae, but is produced in a matrix, and that the larvae contribute to this only by forming and maintaining the matrix. Although there are reports that pig cadavers decomposing without insects may heat up due to microbial activity (21), we compared beetle colonies against meat only and the heat emission was always much larger with the beetles. Therefore, beetle-derived matrix on meat surface may be viewed as a single key factor responsible for the heating effects reported in this study. However, heating of the feeding matrix and the cadaver may superimpose and we suspect that in a natural, large cadaver situation this may lead to an increased total effect.

In the experiments we used fresh pork meat (pieces of 80-150 g), whereas in the field situation *N. littoralis* colonizes large cadavers in a late decomposition stage, with regular occurrence on carcasses larger than 30 kg (13, 56). This may raise doubts as to the translation of our findings to the natural, field situation. When blow flies are absent (in central Europe during early spring), *N. littoralis* may drive active decay of large carrion, replacing blow fly larvae (49, 57). In such cases several hundreds of adult beetles may visit a cadaver and thousands of *N. littoralis* larvae may be present, with minimal or absent colonization by other necrophagous insects, which amounts to the carrion monopolization by these beetles (Fig. 9a, b in ESM). When a cadaver is infested with blow fly larvae and it is large enough not to be entirely consumed by blow flies, the active decay driven by maggots is frequently followed by the active decay driven by larval *N. littoralis* (49). In such cases, *N. littoralis* may be also very abundant and monopolize carrion that is left after blow flies cease feeding (Fig. 9c, d in ESM). Therefore, the monopolization of carrion by *N. littoralis* occurs in a natural, field situation. Moreover, during our field experiments, we regularly observed that *Necrodes* larvae formed a feeding matrix on such monopolized carcasses and were able to spread their exudates over large carrion parts and even the entire surface of a large pig carcass (Fig. 9 in ESM). When larval aggregations of *N. littoralis* were present, they had distinctly higher inner temperature than ambient air (19). Therefore, the formation of feeding matrix by *N. littoralis* and heat production in the matrix are not the laboratory artefacts, but natural phenomena. The mating and oviposition behaviour of adult beetles that were provided with fresh pieces of pork meat were normal; beetles eagerly undertook these actions in our colonies. Moreover, premature development of the *N. littoralis* beetles that were reared on the fresh pork diet to collect developmental data for forensic applications was also normal (58). Although beetles in the field situation feed on a more decomposed carrion, with a more diverse and abundant microbiome (59), using the fresh meat in our experiments cannot explain the differences that were observed between the treatments, as in all the setups fresh pork was used. Experimental design with intact carcasses (e.g. rabbits) would be more ecologically relevant, but also less controllable and more difficult to replicate in the laboratory.

The feeding matrix is a complex microenvironment. Its properties certainly change along the decomposition timeline, with a beetle life stage that participates in its formation or with initial condition of carrion. Beetles contribute to its formation and maintenance by spreading their exudates over carrion surface. The anal exudates of *Necrodes* adult beetles are mixtures of anal secretions with fecal material (54, 55), whereas exudates of larvae are probably dominated by feces. Beetles may also apply oral exudates. Exudates of adult and larval beetles may contain digestive compounds, antimicrobials, beneficial microbes, undigested food etc. Accordingly, beetles may contribute to the matrix formation and maintenance mostly through exodigestion of carrion and manipulation of its microbial content with probable large differences in this respect between species and life stages.

Although our results do not support any of the putative mechanisms for the heat generation in the matrix, two general hypotheses may be formulated in this respect. The heat production may involve the activity of microbes. Feeding matrix on carrion, thoroughly investigated in burying beetles, is a mixture of microbes, their metabolites and compounds released into the matrix directly by the beetles or through carrion digestion (23, 25). Microbes involved in composting produce heat as a by-product of their metabolic processes (60). This mechanism is used by megapode birds and crocodilians to incubate their eggs (6-9). A similar mechanism may be harnessed by *Necrodes* beetles to produce heat on carrion. Heat may however also be produced through exothermal digestion of carrion by beetle exudates. Exothermic digestion has been hypothesized to produce heat in the aggregations of blow fly larvae, although there is no detailed description for this mechanism or evidence in support (17).

The heat of a matrix may benefit *Necrodes* larvae in several ways. By increasing larval development rate, heat may reduce time larvae spend on carrion. Our results support this interpretation. When we compared M+A+L and M+L colonies, we found that heat production in a matrix during the larval feeding phase became larger and larval development times became shorter, following the application of exudates by adult beetles (M+A+L colonies). The colonies differed only in the presence of adult beetles and the application of their exudates on meat in the pre-larval phase. Accordingly, heat and development effects recorded in this experiment may be robustly attributed to these differences. Because, similarly to other insects, *N. littoralis* reveals an increase in development rate and a decrease in development time with an increase in a rearing temperature (58), the larger heat of the matrix must have resulted in shorter development time of the larvae. Linking shorter development with an increase in the matrix heat is the most credible and parsimonious interpretation of our results. Exudates of adult burying beetles inhibit carrion decay, facilitate its digestion and change its microbiome (23, 25). Although these effects have not been reported for *Necrodes* beetles, it is probable that at least some of them occur in this species, and may enhance development, similarly to the matrix heat. From this point of view, shorter larval development may be an effect of the cumulative action of the matrix heat and other matrix-related factors. Interestingly, shorter larval development times were found also for *Nicrophorus mexicanus* (61) and *Nicrophorus vespilloides* (27) in full care or prehatch care conditions (i.e. with exudates application by adult beetles) as compared to the no care conditions, and for *Nicrophorus orbicollis* (62) in the biparental care conditions (i.e. more exudates) as compared to the uniparental care conditions. Although no data exist on the heat production in the feeding matrix formed by burying beetles, it is tempting to hypothesize that these effects, similarly to *Necrodes* beetles, resulted from the lagging thermal benefits of carrion preparation by adult *Nicrophorus* beetles.

Pukowski (24) suggested that smearing carrion with anal exudates may favour aggregation of *Nicrophorus* larvae. Larvae of *N. littoralis* consistently respond to heat in their environment, they aggregate in the hot spots and respond to changes in the hot spot location by relocating themselves (13). Accordingly, heat of the matrix may stimulate larvae to form aggregations in a suitable site for their feeding.

However, the benefits of *Necrodes* larvae may be less obvious. The heat of the matrix may inhibit or support the growth of microbial components of the matrix. Studies of burying beetles linked the microbial suppression on carrion with the production of antimicrobial compounds (23, 25, 28, 37, 38, 40). The heat of the matrix may act synergistically with antimicrobials. There is evidence, from other insect groups, that heat may act in this way. A wax moth larvae *Galleria mellonella* (Lepidoptera: Pyralidae), colonizing dead or weak honeybee colonies, were found to elevate temperature when aggregated inside beehives (5, 63). A recent study showed substantial mortality of *Galleria* larvae after infection with *Metarhizium* fungi at 24°C, whereas at 34°C a 10-fold higher dose of the fungus was necessary to reach similar mortality rate, indicating that temperature elevation by communally feeding *Galleria* larvae suppresses entomopathogenic fungi (64). Consequently, the larger heat production in the matrix formed by larvae after application of exudates by adult *Necrodes* beetles, and the lower mortality in such colonies suggest that the matrix heat facilitates suppression of microbial competitors, and favour beetle survival on carrion. Heat of a matrix may also support growth or activity of mutualistic microbes. Duarte et al. (34) hypothesized that *Nicrophorus* beetles seed their inner microbes on carrion or replant microbes from the carrion gut to its surface. Nevertheless, Miller et al. (65) found little similarity between microbiomes of adult *N. defodiens* gut, their anal secretions and prepared carrion, suggesting that only key microbes are transmitted from the beetle gut to the carcass. Recent analyses of carrion beetle microbiomes recurrently reported an abundant presence of *Yarrowia* yeasts in the beetle gut and on carrion, which suggests a mutualistic link between the yeasts and the beetles (23, 25, 35, 41, 42). Microbial mutualists are probably also associated with *Necrodes* beetles, and the heat of the matrix may support their growth or activity.

Our hypotheses on the benefits of the matrix heat do not exclude themselves. Moreover, heat is not the only attribute of the matrix that is beneficial for the beetles. We assume that microbial and enzymatic content of the matrix brings clear benefits for *Necrodes* larvae, similarly to the *Nicrophorus* larvae. Therefore, our results on the larval mortality most likely resulted from the cumulative action of the heat and the other beneficial attributes of the matrix.

Interestingly, we found that larval mortality increased when we extended the pre-larval phase. The effect was present in M+A+L and M+L colonies. It was very clear in M+A+L colonies, and because adult *Necrodes* beetles were feeding on the meat and we did not supplement it, this must have decreased the quantity of meat available for the larvae and as a consequence their survival. Extension of the pre-larval phase decreases the quality of meat as well, due to the microbial decomposition. Although spreading the exudates of adult beetles probably suppresses putrefaction (36), it can have the inhibitory effect towards decomposition mainly on the meat surface and does not inhibit microbes in the inner layers of meat. Application of exudates by adult beetles has a rather quantitative effect that suppresses but not eliminates putrefaction. Thus, the decreasing quantity and quality of meat with the extension of the pre-larval phase explain this part of our results.

Although scenarios for the evolution of behavioural strategies of spreading the exudates over carrion are highly speculative, several hypotheses may be considered in this respect. Suppression of putrefaction, heat production, facilitating exodigestion and enhancing the growth of beneficial microbes might have exerted selection pressures here. The extent to which these effects were important for different carrion beetles is open to future research, but it will be very difficult to study these effects in isolation. Moreover, their importance certainly differs between the species and also between the life stages of the species. *Necrodes* beetles are communal breeders, abundantly colonizing large cadavers (56, 66), whereas *Nicrophorus* beetles usually breed individually, monopolizing small cadavers (11, 24). Therefore, spreading the exudates may be associated with different constraints and benefits in these beetles.

An increase in heat emission of the matrix formed by larvae after the application of exudates by adult beetles in the pre-larval phase, in association with a decrease of larval development time and mortality, may be regarded as a new form of indirect parental care with an important thermal component. This is also the first demonstration of parental care among Silphinae beetles. Although anal secretions of *Necrodes surinamensis* were found to have antibacterial action (36), no previous study of *Necrodes* beetles reported that they spread the exudates over carrion or that spreading the exudates enhances their offspring’s fitness. Surprisingly, Hoback et al. indicated in their paper that anal secretion’s use “…as a preservative seems unlikely because *N. surinamensis* arrives at large (>500 g) carcasses and its larvae feed on maggots” (36). The other studies on anal secretions of *Necrodes* beetles were focused on their defensive function (54, 55). The hypothesis of parental care of *Necrodes* beetles needs further studies and more data in support, as there are alternative explanations for the behaviour of adult *Necrodes* beetles on carrion. The beetles may simply feed on carrion while engaging in reproduction and through defecation they may increase a microbial load on carrion surface with side effects for their offspring.

Highly developed parental care occurs in *Nicrophorus* beetles (Nicrophorinae), with concealment and preparation of small carrion, provisioning of pre-digested food for young larvae and defence of brood or carrion against competitors (11, 24). Full parental care of *Nicrophorus* beetles results in higher larval survival and mass, compared to broods without parental care (27, 67). Simple parental care, in the form of clearing carrion of fly larvae and brood guarding was also described in *Ptomascopus* beetles (Nicrophorinae) (68, 69). Our findings suggest that indirect forms of parental care, particularly carrion manipulations bringing deferred benefits for the larvae, may be more common among carrion beetles.

Feeding matrixes formed by other insect groups on various substrates may produce heat, equally to the matrix formed on carrion by *Necrodes* beetles. Blow fly larvae (Diptera: Calliphoridae) while feeding on carrion elevate the temperature inside their aggregation (17, 70). Similarly, wax moth larvae *Galleria mellonella* (Lepidoptera: Pyralidae) elevate the temperature when feeding in aggregation inside a beehive (5, 63). Thermogenesis may occur in these cases in the feeding matrix that is formed on carrion or bee chambers by the larvae. Although such thermogenesis is not known in other insect groups that form the matrix, e.g. burying beetles (23), our findings suggest that it may be more frequent, particularly among species that feed on rich and bulky resources, e.g. carrion, fallen fruits or dung.

## Methods

### Beetle colony maintenance and general rearing protocols

Laboratory colony was established using adult beetles sampled from pig carcasses in alder forest of Biedrusko military range (52°31’N, 16°54’E; Western Poland). Beetles were kept in rearing containers (20-30 insects per container, sex ratio about 1:1) on humid soil, at room temperature (20-23°C) and humidity (50-60%). Three to five containers were usually maintained at the same time in the laboratory. Colonies were provided with fresh pork meat *ad libitum* (raw pork in pieces: shoulder, neck or ham).

All experiments were performed in rearing containers with a volume of 1.5 l (initial experiments and experiment 1) or 3.5 l (experiment 2) with about 5 cm of humid soil. The soil was not autoclaved. On one side of container meat was placed, and on the opposite side, we put wet cotton wool to maintain high humidity inside and water for the beetles (Fig. 3). To prevent the meat from drying out and to mimic skin that covers carrion and below which *N. littoralis* usually feed and breed, we put aluminium foil over the entire surface.

*N. littoralis* lays eggs (30-50 per batch) to the soil. There are three larval instars. After the third instar larva ceases feeding, it buries itself in the soil, forms a pupal chamber where it pupates. After emergence, adult beetle stays for some time in the chamber, then it digs itself out fully coloured.

### Thermal imaging and temperature quantification

We monitored thermal conditions inside beetle colonies using thermal imaging camera Testo 885-2 with 30° x 23° infrared lens (Testo, Germany), mounted to a tripod while making images. Images were taken in room temperature and humidity (20-23°C, 50-60%), with rearing containers taken outside of a temperature chamber and images made within no more than 2 minutes.

Based on initial tests with meat, all temperature measurements were made with the emissivity set at 0.8 and reflected temperature set at 17°C. Heat production of the matrix-covered surface was defined as a difference between the average temperature of the surface and the background. To obtain background surface temperatures, we measured average temperature in an area of clean soil located in the meat-opposite side of the container. These measurements were averaged for the first three days starting from the colony establishment. During these initial days, the matrix temperature was usually only a little higher than the background temperature, so the heat production in the matrix had negligible effects on the background temperature. Because the matrix enlarged with colony age, and at some time it covered also the soil that surrounded meat, it was impossible to quantify heat production always within the same area. Instead, we quantified the average temperature in the largest possible circular or ellipsoidal area, depending on the shape of the surface covered with the matrix. We used for this purpose in-built tools of the IRSoft 4.5 software (Testo, Germany).

### Initial experiments

To determine the optimal number of larvae and the optimal quantity of meat, we performed several initial trials. By comparing heat production in larval colonies of various abundance (10, 20, 30, 40, 50 and 100 larvae), we found that normal growth and heat production were already present in colonies of 40 larvae (Fig. 10 in ESM). Accordingly, we used such colonies in the main experiments. During initial trials, we also tested setups with different quantity of meat and decided to use 2 g per larva in Experiment 1 and 3.5 g per larva in Experiment 2. In Experiment 2 we investigated the effect of heat in a feeding matrix on larval fitness, so we decided to provide larvae with more meat to maintain their optimal growth.

### Environmental temperature and heat production in feeding matrix (Experiment 1)

To test the effect of rearing temperature on the heat production in the matrix, we compared heat emission of the matrix across larval colonies reared under constant temperatures of 16, 18, 20 and 23 °C. Heat production in larval colonies (M+L setup, 40 larvae; 80-85 g of raw pork in pieces: shoulder, neck or ham) was compared against heat production in paired containers with meat only (M setup, 80-85 g of pork). Ten pairs of containers (replicates) were studied in each temperature (nine pairs in 20 °C). To establish experimental colonies we sampled freshly hatched first instar larvae from our main colony. Containers were kept in temperature chambers (ST 1/1 BASIC or +, POL EKO, Poland). Two temperatures were studied at the same time. Experiments in 18 and 23 °C started on 28 January 2019 and in 16 and 20 °C on 19 March 2019. To keep thermal conditions as close as possible within the replicate pairs of containers, we kept each pair on the same shelf in the chamber. Once a day colonies were taken out of a chamber to make thermal images. Results were analysed using ANOVA for repeated measures designs in Statistica 13 (TIBCO Software Inc., US), with rearing temperature as an independent variable, container type (M+L or M setups) in a pair as a repeated measures variable and the average heat production in the matrix (i.e. heat production averaged across measurement days) as a dependent variable. No outliers were detected. Fisher LSD test was used for post-hoc pairwise comparisons.

### Heat production and beetle development in different colonies (Experiment 2)

Our main experiment compared thermal conditions and beetle development between two types of colonies in a paired experimental design (Fig. 1). The first colony type had adult beetles in the pre-larval phase and larvae in the larval feeding phase (M+A+L setup). The second type had no adult beetles in the pre-larval phase and larvae in the larval feeding phase (M+L setup). Because we also wanted to test, if the length of the pre-larval phase affects larval fitness, we used three durations of this phase (3, 4 and 5 days). Each treatment was replicated 10 times. Pairs of colonies (M+A+L and M+L setups) were replicates. In each pair, insect colonies were established using larvae hatched at the same time and sampled at random from the same rearing container, meat from the same piece of pork and soil from the same package. Replicate pairs were kept on the same shelf in a temperature chamber at 23 °C (ST 1/1 BASIC or +, POL EKO, Poland). Beetles were provided with 145-150 g of pork (raw pork in pieces from shoulder, neck or ham) in both setups. In the M+A+L setup 10 adult beetles (sampled at random from our main colony, sex ratio 1:1) were kept on meat for the duration of 3, 4 or 5 days, depending on the treatment. Afterwards, the meat was transferred to a new container and 40 freshly hatched first instar larvae were added. Meat relocation was necessary because adult beetles usually oviposited during the pre-larval phase and it was difficult to control the number of larvae in the colonies. In the M+L setup, meat was decomposing without insects for the duration of 3, 4 or 5 days, depending on the treatment. Then, it was transferred to a new container and 40 freshly hatched first instar larvae were added to the container. Experiments started on 3 March 2019 (3 replicates), 15 March 2019 (7 replicates), 8 May 2019 (10 replicates) and 5 August 2019 (10 replicates). Once a day colonies were taken out of the chambers to make thermal images.

After larvae stopped feeding and began to bury themselves, we counted and weighed them (laboratory scale AS 82/220.R2, Radwag, Poland). Then, larvae were transferred to the soil-filled Petri dishes (3 larvae per dish), where we monitored further development to determine pupation and eclosion times. Results were analysed using ANOVA for repeated measures designs in Statistica 13 (TIBCO Software Inc., US), with the duration of the pre-larval phase as an independent variable and a container type (M+A+L or M+L setups) in a pair as a repeated measures variable. The average heat production in the pre-larval phase and the larval feeding phase, the average (per colony) larval and pupal development times, larval mortality and the average (per colony) mass of postfeeding larvae were used as dependent variables in separate analyses. No outliers were detected. Fisher LSD test was used for post-hoc pairwise comparisons.

## Acknowledgments

The study was funded by the National Science Centre of Poland (grant no. 2016/21/B/NZ8/00788).

## Ethics statement

This manuscript describes laboratory experiments using carrion beetle species *N. littoralis*. The species is not under protection. No permission or approval from the Ethics Commission was needed.

## Data accessibility statement

The datasets supporting this article have been uploaded as part of the supplementary material.

## Competing interests’ statement

We have no competing interests to declare.

## Authors’ contributions statement

S.M. developed the concept for the study and the article, analysed the data and wrote the manuscript. Both authors performed experiments, prepared raw data for analyses, discussed the results, prepared figures and reviewed the manuscript.

## Notes

### Competing Interest Statement

The authors have declared no competing interest.

### Summary of Updates

Revised version of the manuscript.

