## supplementary figures for "Heat production in a feeding matrix formed on carrion by communally breeding beetles"

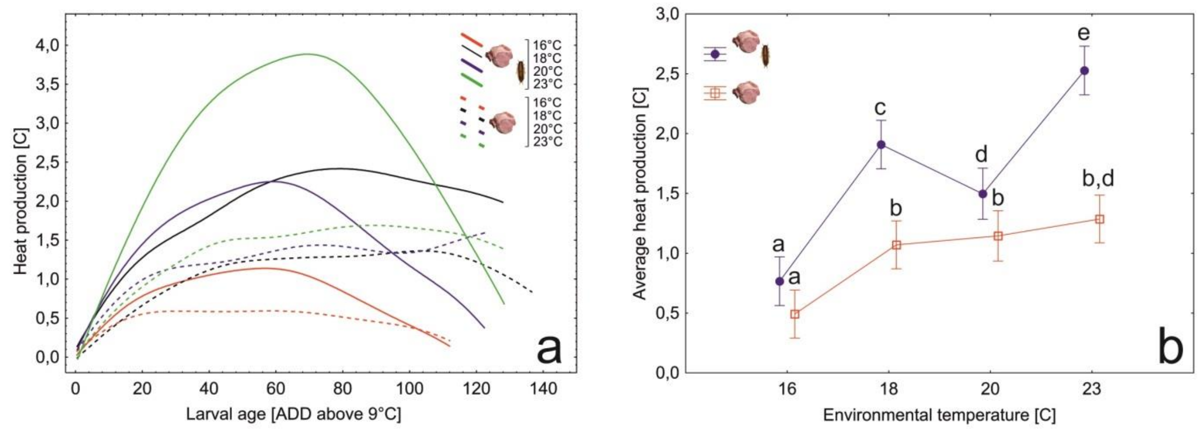


Fig. 5. Longitudinal thermal profiles (a) and differences in the average heat production (b) of the feeding matrix between colonies of larval *Necrodes littoralis* reared under different constant environmental temperatures, compared against thermal profiles and heat production in meat decomposing without the beetles. Thermal profiles were fitted to the data using the distance-weighted least-squares smoothing procedure. ADD – accumulated degree-days, symbols – means, whiskers – 95% confidence intervals, different letters denote significant differences in pairwise comparisons.

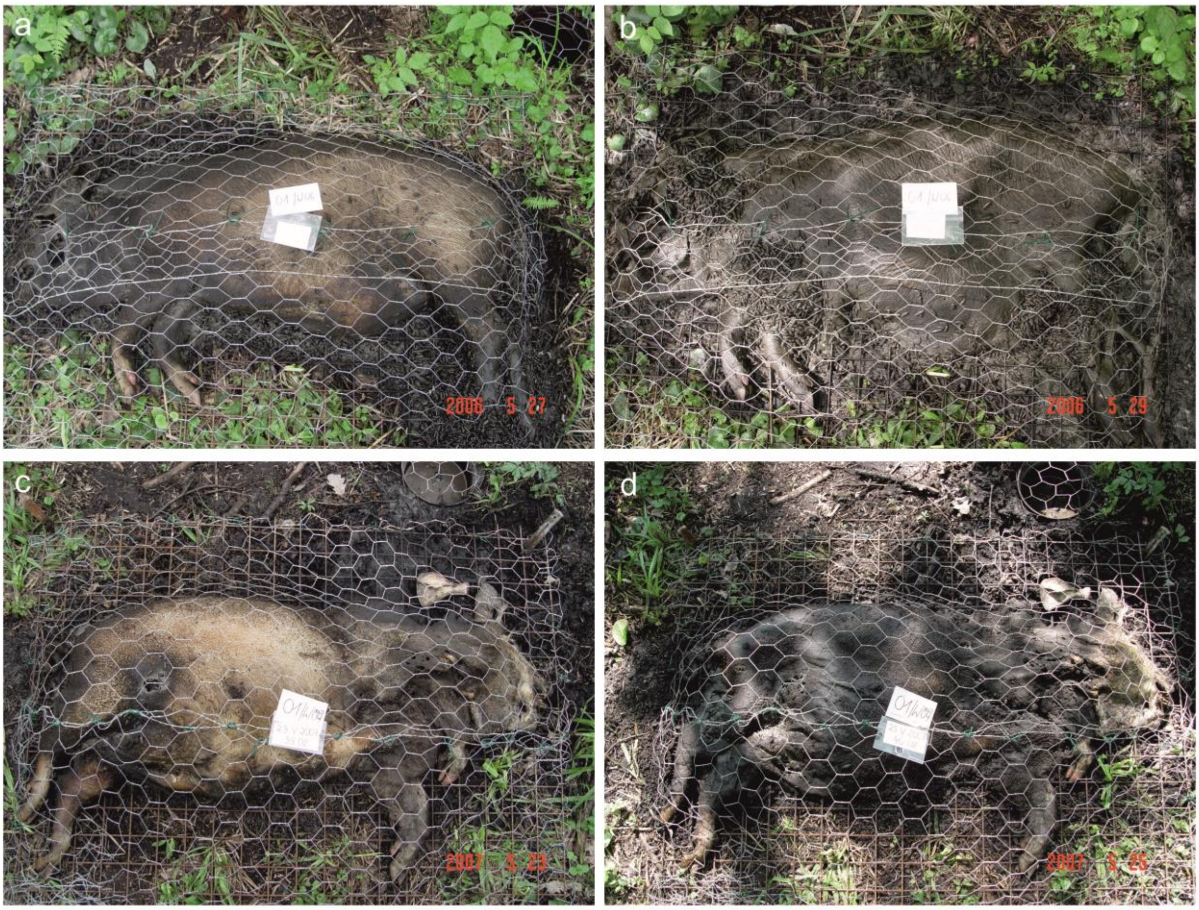


Fig. 9. Monopolization of pig carcasses by *Necrodes littoralis.* a,b – Carcass completely monopolized by larvae of *N. littoralis*, a – 34 day of decomposition, larval masses of *N. littoralis* visible near the hind legs, larval exudates cover large parts of the cadaver, b – 36 day of decomposition, larval masses visible near the legs (fore and hind) and the neck, exudates cover the entire cadaver. c,d - Carcass monopolized in part by larvae of *N. littoralis*, the anterior part of the cadaver was consumed by larval blow fies, c – 38 day of decomposition, exudates of larval *N. littoralis* cover parts of the trunk and hind legs, d – 40 day of decomposition, larval exudates cover the entire trunk and hind legs.

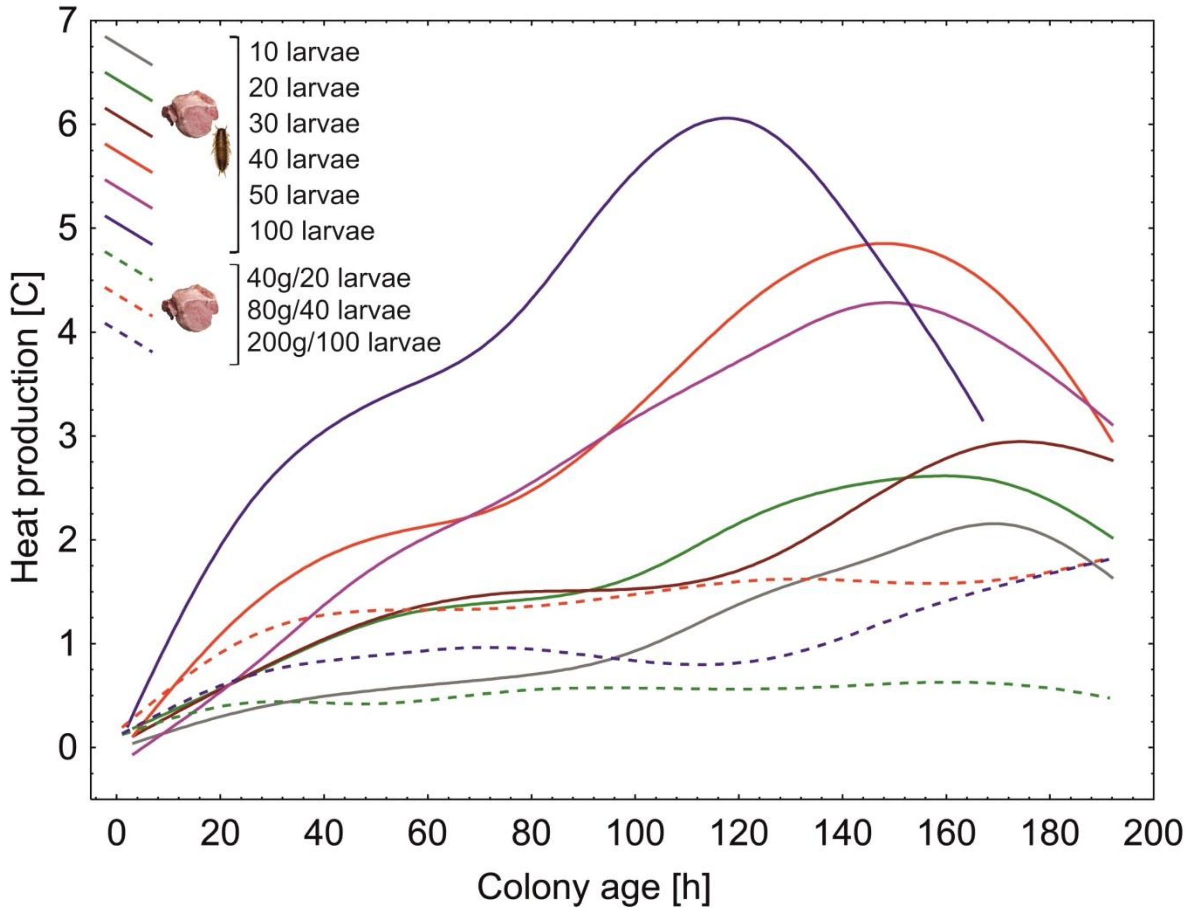


Fig. 10. Longitudinal thermal profiles of the feeding matrix in colonies of larval *Necrodes littoralis* with different numbers of larvae (solid lines) and control colonies with meat only (dashed lines). Colonies were reared at room temperature (20-23°C). Thermal profiles were fitted to the data using the distance-weighted least-squares smoothing procedure.
